# Calcium negatively regulates secretion from dense granules in *Toxoplasma gondii*

**DOI:** 10.1101/386722

**Authors:** Nicholas J Katris, Geoffrey I McFadden, Giel G. van Dooren, Ross F Waller

## Abstract

Apicomplexan parasites including *Toxoplasma gondii* and *Plasmodium* spp. manufacture a complex arsenal of secreted proteins used to interact with and manipulate their host environment. These proteins are organised into three principle exocytotic compartment types according to their functions: micronemes for extracellular attachment and motility, rhoptries for host cell penetration, and dense granules for subsequent manipulation of the host intracellular environment. The order and timing of these events during the parasite’s invasion cycle dictates when exocytosis from each compartment occurs. Tight control of compartment secretion is, therefore, an integral part of apicomplexan biology. Control of microneme exocytosis is best understood, where cytosolic intermediate molecular messengers cGMP and Ca^2+^ act as positive signals. The mechanisms for controlling secretion from rhoptries and dense granules, however, are virtually unknown. Here, we present evidence that dense granule exocytosis is negatively regulated by cytosolic Ca^2+^, and we show that this Ca^2+^-mediated response is contingent on the function of calcium-dependent protein kinases *Tg*CDPK1 and *Tg*CDPK3. Reciprocal control of micronemes and dense granules provides an elegant solution to the mutually exclusive functions of these exocytotic compartments in parasite invasion cycles and further demonstrates the central role that Ca^2+^ signalling plays in the invasion biology of apicomplexan parasites.

## Introduction

Apicomplexan parasites comprise a large phylum of primarily obligate intracellular parasites of humans and animals that have a significant impact on human health and livestock production. Notable apicomplexans genera include: blood parasites *Plasmodium* (causative agents of malaria), *Babesia* and *Theileria* (common cattle parasites); enteric epithelial parasites *Cryptosporidium* and *Eimeria*; and systemic parasites *Toxoplasma* and *Neospora*. The phylum embraces at least 6,000 species with global distribution infecting animals and even other protists (Adl et al., 2012).

Moreover, metagenomic environmental sampling shows that apicomplexans can be dominant components of natural communities indicating significant roles in ecosystems and the evolutionary success of this group (de Vargas et al., 2015; Mahé et al., 2017). One key to the success of apicomplexans is their efficient infection cycles in which they select their host cell, penetrate it non-destructively, feed and multiply within this cell while subduing or deflecting host organism defences, and finally escape from the host cell releasing multiple progeny (Blader, Coleman, Chen, & Gubbels, 2015). This cycle is largely mediated by co-ordinated release of a number of different exocytotic compartments delivering cargo to a range of extracellular destinations.

*Toxoplasma gondii* has served as a model for apicomplexan infection cycle events with three categories of secretory compartments identified—micronemes, rhoptries and dense granules—that facilitate the major events of the invasion cycle (Carruthers & Sibley, 1997). Micronemes are exocytosed when the parasite is searching for a host cell, and secreted microneme proteins (MICs) decorate the parasite cell surface to act as attachment ligands and enable the characteristic gliding motility of the group (Frénal, Dubremetz, Lebrun, & Soldati-Favre, 2017). Upon selection of a cell to invade, proteins from rhoptry organelles are then secreted into the host, forming a ‘moving junction’ entry structure through which the parasite penetrates the host (Guérin et al., 2017). As the parasite enters, a host plasma membrane-derived parasitophorous vacuole (PV) invaginates and surrounds the parasite. PV formation is accompanied by secretion of further rhoptry proteins into the host, some of which actively block host attack of this new internal foreign body (Etheridge et al., 2014; Håkansson, Charron, & Sibley, 2001). Completion of invasion isolates the PV from the plasma membrane, and a third wave of secretion from the dense granules now occurs (Carruthers & Sibley, 1997; Dubremetz, Achbarou, Bermudes, & Joiner, 1993; Mercier & Cesbron-Delauw, 2015; Sibley, Niesman, Parmley, & Cesbron-Delauw, 1995). Dense granule proteins (GRAs) populate and modify the PV membrane for nutrient uptake, and help create an elaborate PV-contained membranous nanotubular network (MNN) (Mercier, Adjogble, Däubener, & Delauw, 2005; Sibley et al., 1995).

Other GRAs target the host cytoplasm and nucleus, and actively reprogram host cell regulatory pathways and functions to facilitate parasite survival and growth (Hakimi, Olias, & Sibley, 2017). After multiple rounds of parasite division, a new infection cycle begins with the secretion of MICs that disrupt host membranes and reactivate gliding motility for escape, dissemination, and targeting of new host cells (Kafsack et al., 2009). Broadly, control of secretion from micronemes is critical for the extracellular stages of the *Toxoplasma* infection cycle, control of rhoptry release for the invasion events, and control of dense granule release for the establishment and maintenance of the host cell environment for the parasite. The coordination of organelle-specific exocytosis is, therefore, a central feature of the parasite’s biology.

Only the control of microneme exocytosis has been studied and illuminated in any detail. The elevation of cytosolic calcium ion (Ca^2+^) levels by release from intracellular stores signals release of MICs to the extracellular environment (Carruthers, Giddings, & Sibley, 1999a; Sidik et al., 2016). Ca^2+^ also stimulates other processes, including extrusion of the conoid and activation of motility, so Ca^2+^ signalling is clearly part of a broader signalling network of the extracellular events of the invasion cycle (Billker, Lourido, & Sibley, 2009; Borges-Pereira et al., 2015; Graindorge et al., 2016; Stewart et al., 2017; Q. Tang et al., 2014; Wetzel, Chen, Ruiz, Moreno, & Sibley, 2004). Two Ca^2+^-dependent protein kinases, *Tg*CDPK1 and *Tg*CDPK3, are major controllers of Ca^2+^-dependent extracellular processes including MIC secretion (Lourido, Jeschke, Turk, & Sibley, 2013; Lourido et al., 2010; Lourido, Tang, & Sibley, 2012; McCoy, Whitehead, van Dooren, & Tonkin, 2012; Treeck et al., 2014). Loss of function of either results in changes to Ca^2+^-induced microneme exocytosis, although changes are not identical suggesting some level of specialisation and/or cooperativity of these kinases (Lourido et al., 2012). Numerous protein substrates have been identified for both *Tg*CDPK1 and *Tg*CDPK3, further evidence for an elaborate signalling network that they control (Lourido et al., 2013; Treeck et al., 2014). Ca^2+^ also has a downstream and direct role for MIC release with exocytosis of micronemes at the apical plasma membrane facilitated by DOC2.1 that recruits the membrane fusion machinery in a Ca^2+^-dependent manner (Farrell et al., 2012).

Other signalling molecules and stimuli occur upstream of Ca^2+^ and illustrate an even broader network of control processes for parasite sensing of cues for its invasion cycle (Carruthers, Moreno, & Sibley, 1999b). Cyclic guanosine monophosphate (cGMP) activates protein kinase G (PKG), and PKG in turn triggers cytosolic Ca^2+^ flux in *Toxoplasma* and *Plasmodium* (Brochet et al., 2014; Sidik et al., 2016; Stewart et al., 2017). In *P. berghei* PKG acts upon phosphoinositide metabolism that ultimately releases inositol (1,4,5)-trisphosphate (IP3) from diacylglycerol (DAG), and IP3 releases Ca^2+^ stores in many systems (Brochet et al., 2014; Schlossmann et al., 2000). This phospholipid metabolism, notably DAG to phosphatidic acid (PA) interchange, has also been shown to contribute directly to microneme docking at the plasma membrane in *Toxoplasma*, further implicating cGMP-controlled events in MIC release (Bullen et al., 2016). The ultimate stimulation mechanism(s) for these cGMP- and Ca^2+^-dependent events has not been identified, however in *Toxoplasma* a role for changes in potassium ion levels—a relative decrease from intracellular to extracellular environments—appears to have an important role as a cue (Endo, Tokuda, Yagita, & Koyama, 1987). Upon successful entry of parasites into their host cell, all of this activation for MIC release and motility must then be supressed in order for the rhoptry- and dense granule-mediated intracellular events to progress. Recent work suggests that cAMP-signalling that activates protein kinase A (PKA) is involved in reducing cytosolic Ca^2+^ levels upon host cell entry and, in turn, reversing the processes that led to MIC secretion (Jia et al., 2017; Uboldi et al., 2018).

While Ca^2+^ and cGMP have emerged as central signals for positive control of microneme exocytosis, almost nothing is known about how secretion from rhoptries and dense granules is controlled.

Dense granules present a particular conundrum, for while some secretion through these organelles has been characterised as unregulated or constitutive, a strong burst of GRA secretion occurs as a post-invasion event, implying some mechanism for its control (Carruthers & Sibley, 1997; Chaturvedi et al., 1999; Coppens, Andries, Liu, & Cesbron-Delauw, 1999; Dubremetz et al., 1993; Mercier & Cesbron-Delauw, 2015; Sibley et al., 1995). Nevertheless, GRA secretion from extracellular parasites is detectable and has often been used as a presumed invariant secretion control in assays of regulated MIC release. This assumption of GRA behaviour, however, has never been thoroughly tested and, in fact, a decrease in GRA secretion has been seen (but rarely commented upon) when extracellular parasites are treated with some Ca^2+^ agonists (Carruthers et al., 1999b; Farrell et al., 2012; Kafsack et al., 2009; Paul et al., 2015).

Here, we have examined the role of Ca^2+^ and cGMP in the regulation of GRA secretion in extracellular tachyzoites using a range of modulators of both Ca^2+^ and cGMP levels. We have also tested for GRA secretion behaviour in mutant cell lines of *Tg*CDPK1, *Tg*CDPK3 and *Tg*RNG2, and using kinase inhibitors, all of which have known defects in Ca^2+^-or cGMP-dependent secretion (Katris et al., 2014; Lourido et al., 2012; McCoy et al., 2012). Our data consistently indicate that Ca^2+^ has a role in negatively controlling GRA secretion, providing a reciprocal control mechanism to that of MIC secretion in extracellular parasites.

## Results

### Agonists and antagonists of cytosolic Ca^2+^ inversely modulate microneme and dense granule exocytotis

To test for any Ca^2+^-dependent responses in dense granule exocytosis from extracellular tachyzoites, we applied a range of concentrations of two commonly used Ca^2+^ ionophores, ionomycin and A23187, that induce a range in levels of MIC secretion response. A gradual increase in MIC2 secretion (above constitutive levels) was observed with increasing concentrations of both ionomycin and A23187 from 1 to 5 µM (Figure 1A). We simultaneously assayed for secretion of the dense granule proteins GRA1, GRA2 and GRA5. In all cases we saw an inverse response to the ionophore treatment, with decreased secretion of dense granule proteins observed with increasing ionophore concentration (Figure 1A). We also assayed for MIC5 secretion, a protein that is proteolytically processed before sorting to the micronemes. The shorter, processed form is secreted from the micronemes, while the longer pro-form of the protein (proMIC5) is believed to take an alternative route and its secretion has been described as constitutive (Brydges, Harper, Parussini, Coppens, & Carruthers, 2008). We saw reciprocal responses of MIC5 and proMIC5 secretion with Ca^2+^ ionophore treatment: MIC5 release responded positively to Ca^2+^, as for MIC2; while proMIC5 showed reduced secretion with Ca^2+^, similar to the GRA protein responses (Figure 1A).

**Figure 1:**
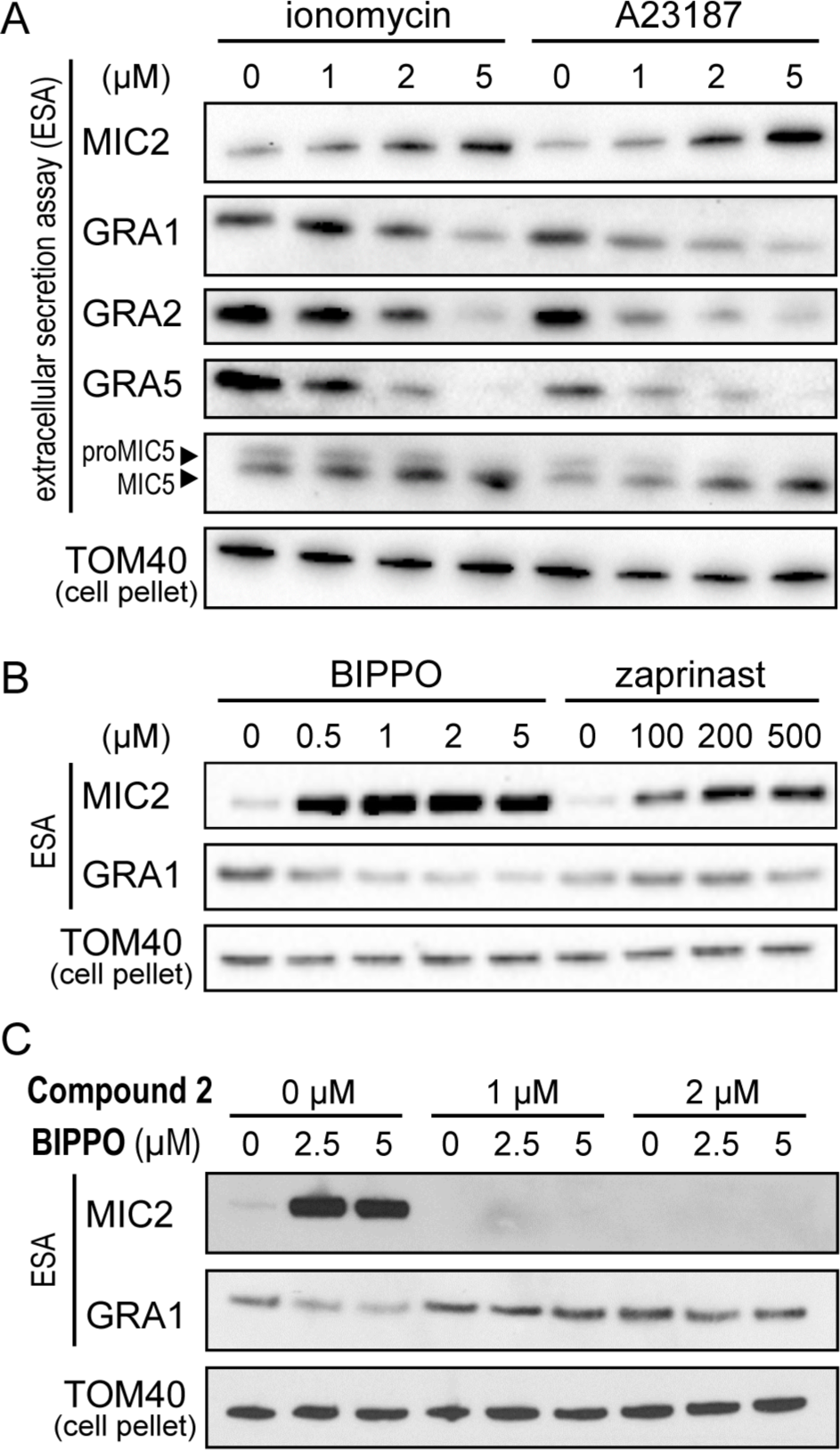
Microneme and dense granule secretion responses to Ca^2+^- and cGMP-based stimulation. Secreted proteins assayed by Western blot of select MICs and GRAs in response to Ca^2+^ ionophores ionomycin and A23187 (A) and phosphodiesterase inhibitors BIPPO and zaprinast (B). (C) PKG inhibitor compound 2 blocks the responses to BIPPO. Mitochondrial protein TOM40 in parasite pellets serve as parasite equivalent loading controls for the extracellular secretion assays (ESA).

Cytosolic Ca^2+^ levels can also be indirectly elevated by activating the PKG signalling pathway with cGMP (Sidik et al., 2016; Stewart et al., 2017). Two phosphodiesterase inhibitors have been widely used to increase cGMP levels: zaprinast, and a more potent analogue 5-benzyl-3-iso-propyl-1H-pyrazolo[4.3-d]pyrimindin-7(6H)-one (BIPPO) (Howard et al., 2015). Increasing concentrations of BIPPO and zaprinast over a range shown to stimulate MIC secretion (0-5 µM and 0-500 µM, respectively) were applied to tachyzoites. MIC2 secretion was far more responsive to BIPPO than zaprinast, with approximately equivalent MIC2 secretion seen with 0.5 µM BIPPO and 200-500 µM zaprinast (Figure 1B). At these concentrations only minor reduction in GRA1 secretion was seen, however, as BIPPO concentrations were further increased through 1 - 5 µM a clear decrease in secreted GRA1 was observed (Figure 1B). To validate that BIPPO was acting through PKG to inhibit dense granule secretion, the PKG inhibitor ‘compound 2’ was tested to see if it could block the secretion responses of BIPPO (Brochet et al., 2014; Donald et al., 2002; Jia et al., 2017; Wiersma et al., 2004). When tachyzoites were pre-treated with compound 2, all constitutive and BIPPO-inducible MIC2 secretion was lost, as well as the BIPPO-induced inhibition of dense granule secretion (Figure 1C).

We further tested for the effect on GRA secretion of modulators of cytosolic Ca^2+^ by treating cells with either BAPTA-AM or thapsigargin. BAPTA-AM is a membrane-permeable Ca^2+^-chelator so treatment reduces available Ca^2+^, whereas thapsigargin is an inhibitor of the sarco/endoplasmic reticulum Ca^2+^ATPase (SERCA) that is believed responsible for recharging sequestered Ca^2+^ pools. Thapsigargin-treatment, thus, leads to cytosolic accumulation of Ca^2+^. BAPTA-AM-treatment resulted in loss of constitutive secretion from micronemes, and no change to dense granule protein secretion, consistent with a role of elevated Ca^2+^ in both of these processes (Figure 2A).

**Figure 2:**
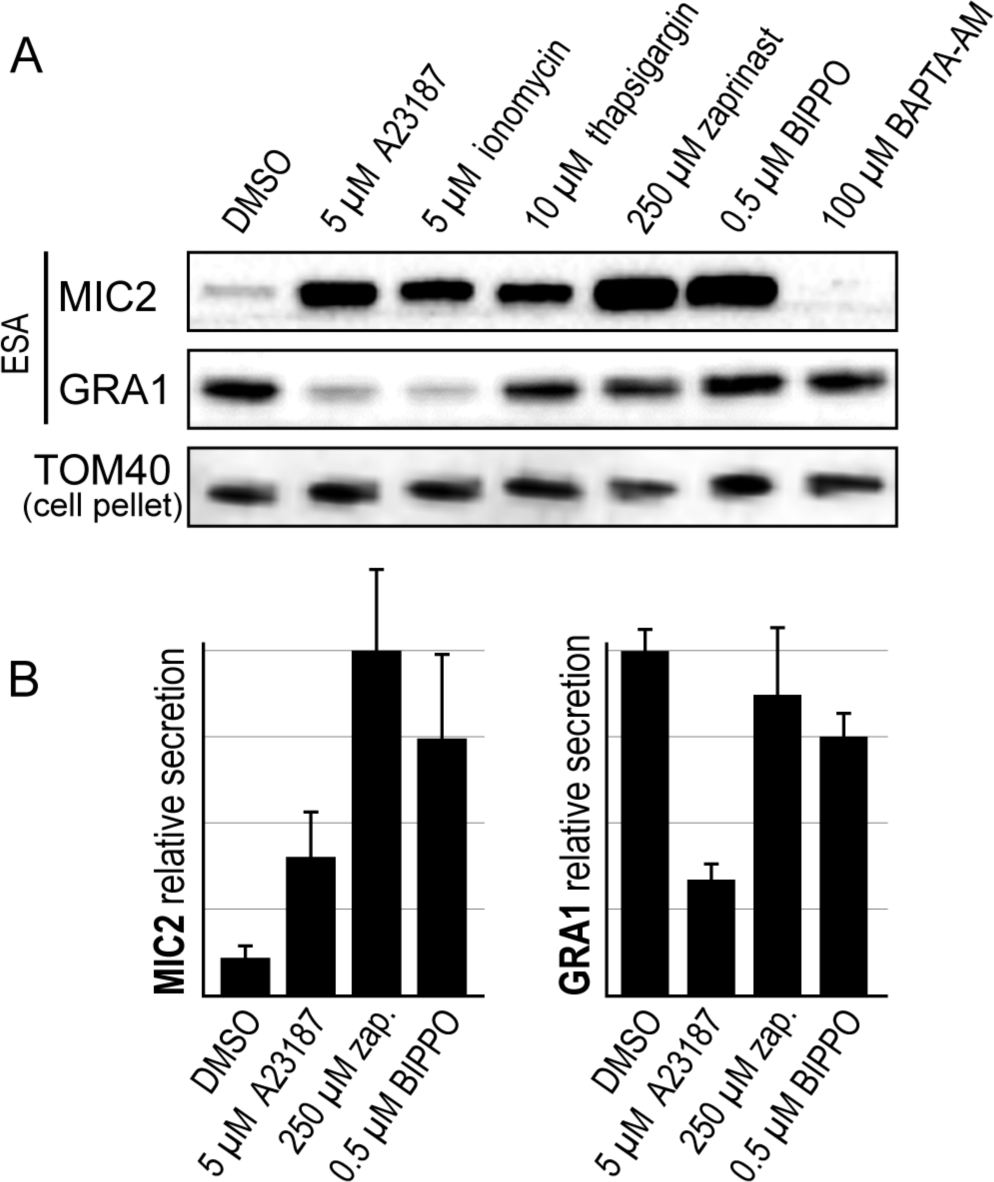
Relative effects of modulators of cytosolic Ca^2+^ on MIC and GRA secretion. (A) Ca^2+^ ionophores A23187 and ionomycin; Ca^2+^ sequestration inhibitor thapsigargin; cGMP agonists zaprinast (zap.) and BIPPO; and Ca^2+^ chelator BAPTA-AM; all affect MIC secretion and have varying effects on GRA secretion. (B) Relative secretion of MIC2 and GRA1 shown over 8 biological replicates. ESA; Extracellular Secretion Assay: TOM40 used as a cell equivalents loading control. Error bars = SEM.

Thapsigargin-treatment resulted in elevated MIC secretion, but no change in GRA secretion when applied at 10 µM (Figure 2A).

In summary, we observe an inverse correlation between MIC and GRA secretion in response to changes in cytosolic Ca^2+^. Further, there is evidence of separation between the manner of eliciting Ca^2+^-signalling and the proportional responses of MIC and GRA secretion. Ionophore treatment to directly release Ca^2+^ stores result in marked increase in MIC secretion and concomitant decrease in GRA secretion (Figure 1 and 2). Conversely, indirect methods for elevating cytosolic Ca^2+^—moderate cGMP stimulus and thapsigargin—result in strong MIC secretion, but proportionately less inhibition of GRA secretion (Figure 1 and 2).

### Mutants in Ca^2+^ signalling disrupt both MIC and GRA secretion responses

Ca^2+^-dependent protein kinases (*Tg*CDPKs) 1 and 3 in *Toxoplasma* are involved in controlling cell processes relevant to invasion and egress, including microneme exocytosis, where elevated Ca^2+^ triggers activation of these processes (Lourido et al., 2010; 2012; McCoy et al., 2012). Given our data that GRA secretion negatively correlates with Ca^2+^ level increase, we tested if *Tg*CDPK1 and/or 3 might be involved in regulating dense granule exocytosis. To test for a role of *Tg*CDPK1 we used an inducible knockdown cell line (iΔHA-*Tg*CDPK1) (Lourido et al., 2010) in which *Tg*CDPK1 levels were strongly depleted after 72 hours of anhydrotetracycline (ATc) treatment (Figure 3A). Untreated (-ATc) iΔHA-*Tg*CDPK1 cells showed typical constitutive MIC and GRA secretion, and both A23187-responsive MIC secretion and coincident inhibition of GRA secretion (Figure 3A-C), consistent with wildtype cells (Figures 1, 2). When *Tg*CDPK1 was depleted (+ATc) levels of MIC secretion were strongly reduced in both constitutive and A23187-treated states compared to the minus-ATc controls. The suppression of GRA secretion with A23187 treatment was also reduced in the *Tg*CDPK1-depleted cells (Figure 3A,C). Therefore, depletion of *Tg*CDPK1 simultaneously results in reductions in both the Ca^2+^-induced increase in microneme exocytosis and inhibition of dense granule exocytosis.

**Figure 3:**
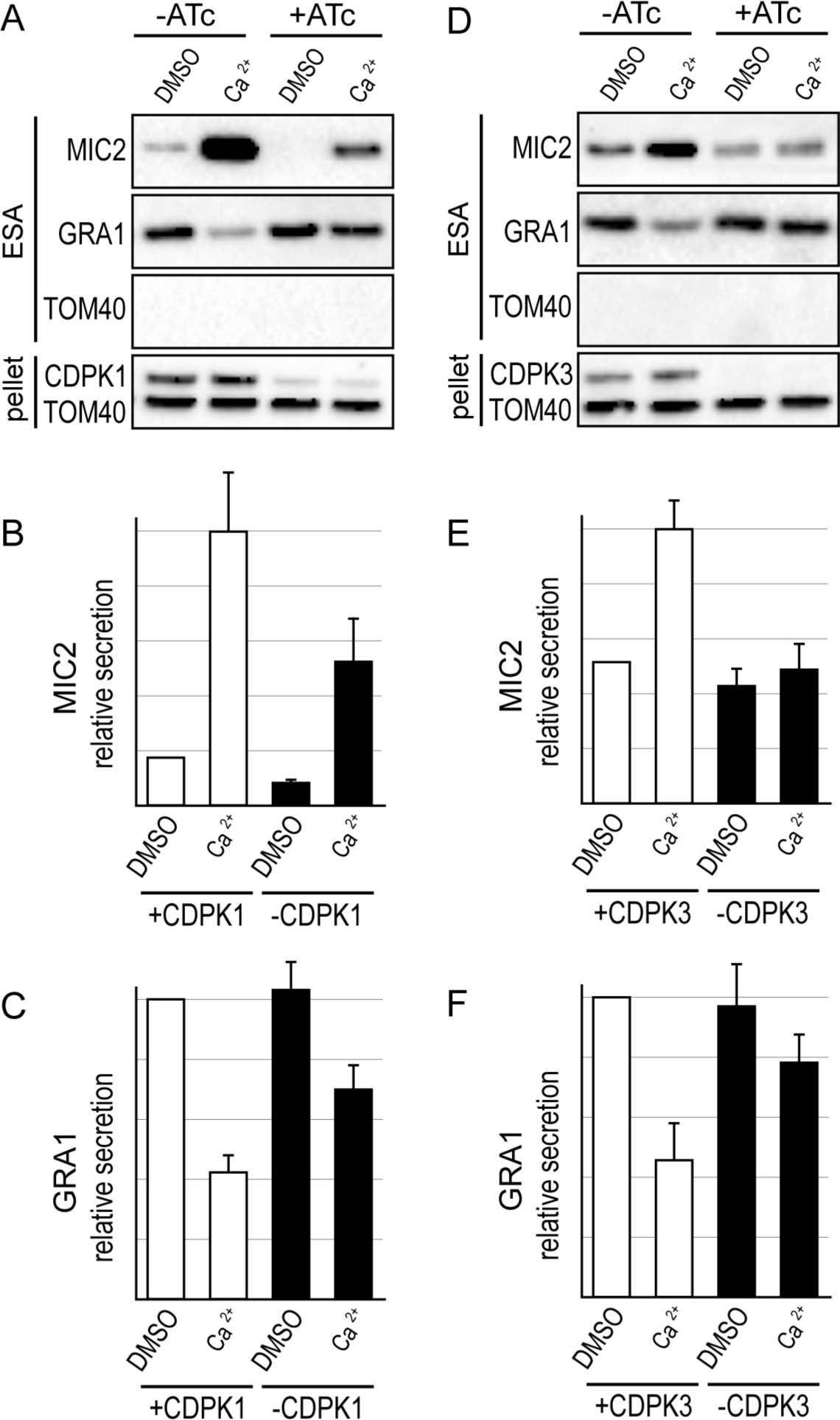
Effect of loss of CDPK1 or CDPK3 on Ca^2+^-induced MIC and GRA secretion. (A-C) Extracellular secretion assays (ESA) with and without ATc-induced CDPK1 depletion in iΔHA-CDPK1 cells. Depletion of HA-CDPK1 is seen by immuno-detection of HA in the cell pellet. A23187 (5 µM) is used for Ca^2+^ stimulation. Relative secretion of MIC2 (B) and GRA1 (C) is shown (n=8). (D-F) ESA and relative secretion (n=7) measurements for wildtype (+CDPK3) versus CDPK3 knockout (-CDPK3) cells. Absence of CDPK3 is seen by CDPK3 immuno-detection in the cell pellet (D). TOM40 in ESA serves a control for cell lysis, and controls for cell equivalent loading in the pellet. Error bars = SEM.

We also used a chemical inhibition strategy to test for the role of *Tg*CDPK1 in the control of secretion from dense granules. The kinase inhibitor 3-methyl-benzyl pyrazolo [3,4-d] pyrimidine (3-MB-PP1) is specific to *Tg*CDPK1 in *T. gondii* (Lourido et al., 2010; 2012), and cells treated with this inhibitor were concurrently assayed for changes of microneme and dense granule exocytosis.

Extracellular parasites were treated with 3-MB-PP1 for 5 minutes post harvesting, and then assayed for protein secretion. Ca^2+^-induced (A23187 treatment) MIC secretion was lost with 3-MB-PP1 treatment, consistent with inhibition of *Tg*CDPK1 (Figure 4Ai). Interestingly, constitutive levels of microneme secretion were unaffected, unlike the *Tg*CDPK1 KD. In these experimental conditions, dense granule exocytosis was also found to be unaffected in that increasing [Ca^2+^] could inhibit levels of GRA secretion, equivalent to the 3-MB-PP1-untreated cells (Figure 4Ai, Bii). Therefore, extracellular parasites showed some behaviours similar to the *Tg*CDPK1 knockdown but not all when treated with 3-MB-PP1 after cell egress. It is possible that *Tg*CDPK1 had phosphorylated targets upon egress, but before 3-MB-PP1 treatment, which might be responsible for the Ca^2+^-inducible dense granule control. To test this, we pre-treated intracellular parasites with the 3-MB-PP1 kinase inhibitor for 5 minutes before mechanical egress and secretion assays. With this pre-harvest treatment, all microneme secretion was lost: both constitutive and Ca^2+^-induced MIC secretion (Figure 4Aii). Furthermore, Ca^2+^-induced inhibition of dense granule protein secretion was also substantially lost (Figure 4Aii, Biii). These results mimic the effects of *Tg*CDPK1 depletion and suggest that in the first 3-MB-PP1 experiment some *Tg*CDPK1 targets were phosphorylated after egress and persisted in this state despite subsequent 3-MB-PP1 inhibition.

**Figure 4:**
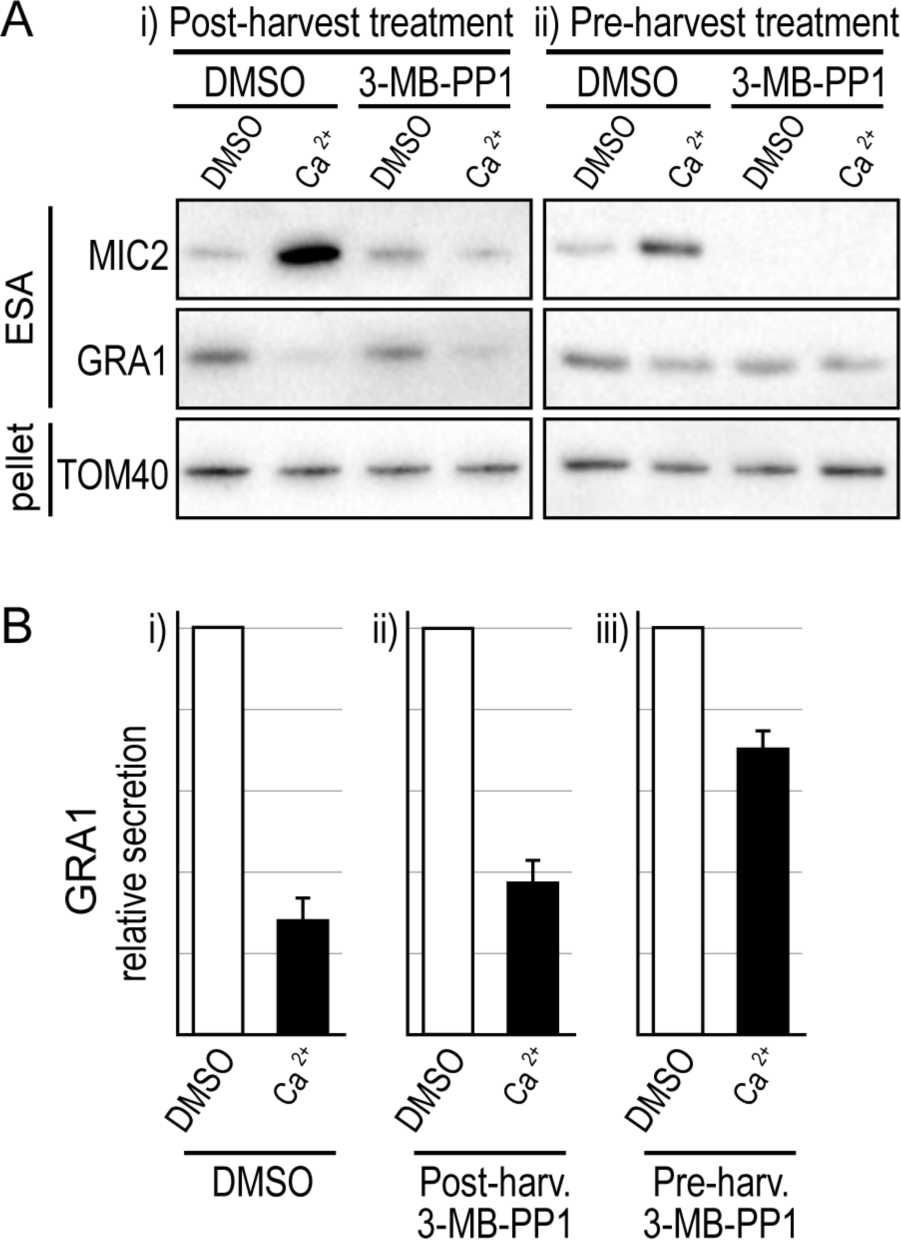
Effect of CDPK1-inhibitor 3-MB-PP1 on Ca^2+^-induced MIC and GRA secretion. (A) Parasites were treated with 3-MB-PP1 either (i) after mechanical egress from host cells, or (ii) before egress, and then Extracellular Secretion Assays (ESA) of MIC2 and GRA1 without or with A23187 (5 µM)-induced Ca^2+^ release. (B) Relative GRA1 secretion was measured for DMSO controls (n=4), post-harvest 3-MB-PP1 (n=4) and pre-harvest 3-MB-PP1 (n=3). Error bars = SEM.

We also tested for a role of *Tg*CDPK3 in GRA secretion regulation using a cell line with the *cdpk3* gene knocked out (Δ*Tg*CDPK3) (McCoy et al., 2012). Unlike the *Tg*CDPK1 KD, *Tg*CDPK3 absence did not affect constitutive MIC secretion (Figure 3D, E). When treated with A23187, Δ *Tg*CDPK3 cells showed neither an increase in MIC secretion, nor a decrease in GRA secretion, suggesting that *Tg*CDPK3 is required for Ca^2+^-mediated control of both of these processes (Figure 3 D-F).

*Tg*RNG2 is an apical complex protein involved in relaying a cGMP signal through to MIC secretion (Katris et al., 2014). Depletion of *Tg*RNG2 interrupts the relay of this signal, although microneme secretion can be rescued with direct Ca^2+^ stimulation by A23187 (Figure 5Ai, Bi). These data suggest that *Tg*RNG2 operates between cGMP sensing and cytosolic Ca^2+^ elevation. We therefore used a *Tg*RNG2 inducible knockdown cell line (iΔHA-*Tg*RNG2) to independently test if dense granule exocytosis control is regulated directly by Ca^2+^ rather than cGMP. *Tg*RNG2-depleted cells showed normal levels of Ca^2+^-induced dense granule exocytosis inhibition, and concurrent elevation of microneme secretion (Figure 5Ai, Bi, Ci). When *Tg*RNG2-depleted cells were activated via cGMP (2.5µM BIPPO), no change in dense granule secretion was seen compared to untreated cells (Figure 5Aii, Cii). These data are consistent with dense granule regulation responding directly to Ca^2+^ and not cGMP.

**Figure 5:**
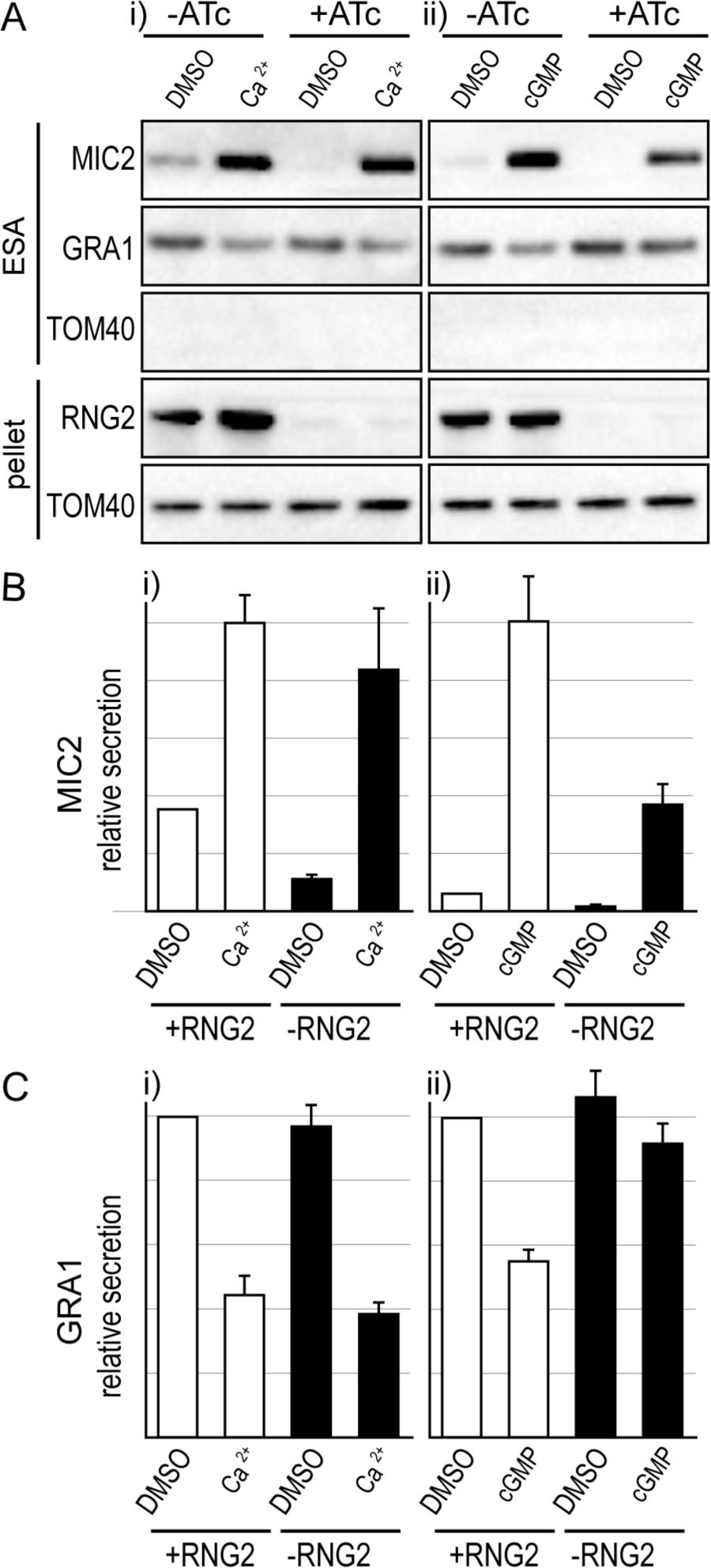
Effect of loss of RNG2 on Ca^2+^- and cGMP-induced MIC and GRA secretion. (A-C) Extracellular secretion assays (ESA) with and without ATc-induced RNG2 depletion in iΔHA-RNG2 cells. Depletion of HA-RNG2 is seen by immuno-detection of HA in the cell pellet. A23187 (5 µM) is used for Ca^2+^ stimulation (i) and BIPPO (2.5 µM) is used for cGMP stimulation (ii). Relative MIC2 (B) and GRA1 (C) is shown for 8 (Ca^2+^, Bi, Ci) and 10 (cGMP, Bii, Cii) biological replicates. TOM40 in ESA serves a control for cell lysis, and controls for cell-equivalents loading in the pellet. Error bars = SEM.

## Discussion

We have tested for changes to rates of dense granule protein (GRA) secretion from extracellular tachyzoites in response to stimuli known to elicit changes in microneme exocytosis, namely, treatments that elevate cytosolic Ca^2+^ (summarised in Figure 6). We consistently see evidence of reduced GRA secretion in conditions that raise cytosolic Ca^2+^, including both ionophore treatment that allows discharge of Ca^2+^ stores into the cytoplasm, and cGMP treatment that indirectly raises cytosolic Ca^2+^ through activation of PKG (Brochet et al., 2014; Sidik et al., 2016; Stewart et al., 2017). Both treatment types increased microneme protein secretion. Mutant cell lines for *Tg*CDPK1, *Tg*CDPK3 and *Tg*RNG2 with known phenotypes in microneme secretion control (Katris et al., 2014; Lourido et al., 2010; McCoy et al., 2012), all similarly displayed reciprocal GRA control phenotypes, further supporting a role for cytosolic Ca^2+^ levels in down regulation of GRA secretion. We also note that several published reports show evidence of Ca^2+^-mediated suppression of secretion of dense granule proteins, although in these cases the effects seen were either not commented on or explored (Carruthers et al., 1999b; Farrell et al., 2012; Kafsack et al., 2009; Paul et al., 2015).

**Figure 6.**
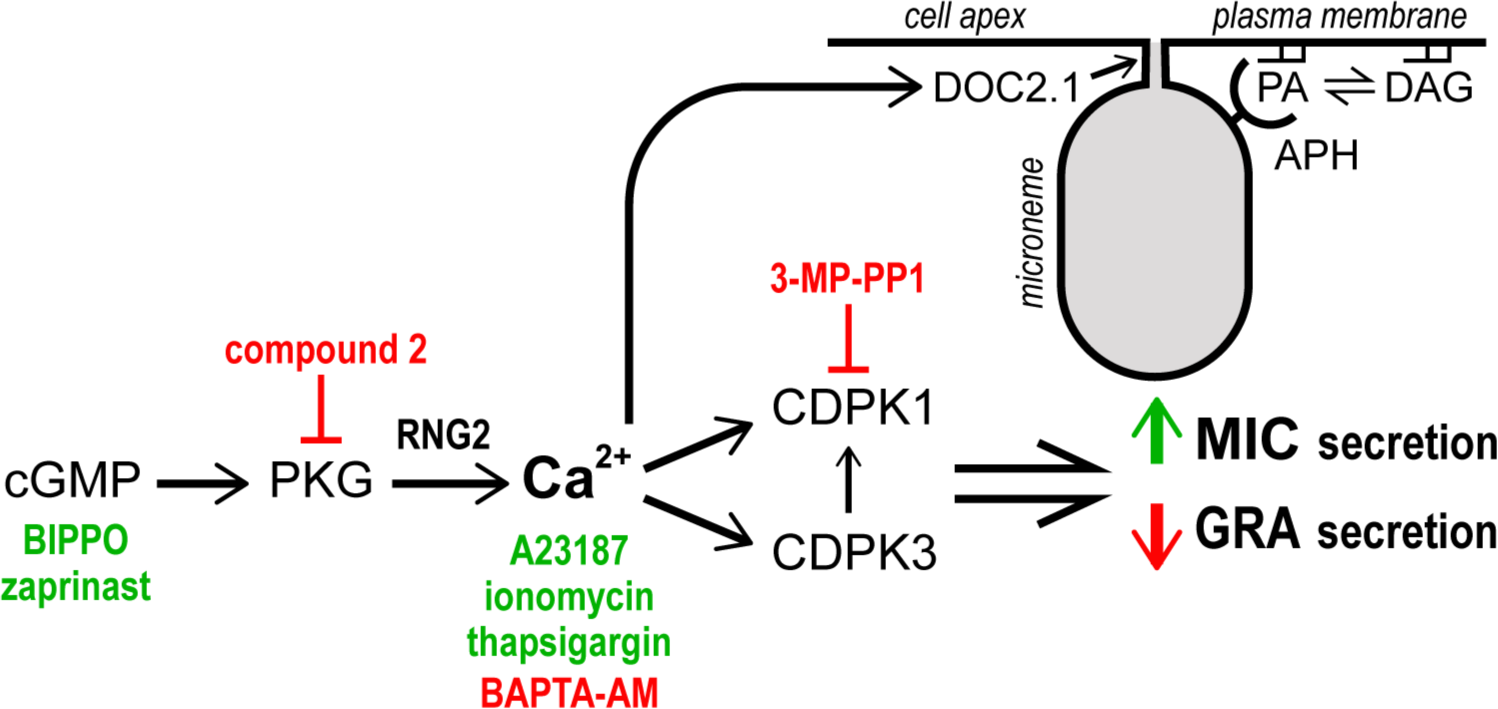
Summary of known regulatory events that contribute to the control of microneme and dense granule exocytosis. Chemical agonists (green) of secondary messengers cGMP and Ca^2+^ and antagonists (red) of these messengers or kinases used in this study are shown. Microneme interaction and fusion at the plasma membrane via APH-PA interactions and Ca^2+^-dependent DOC2.1 activity is also shown. APH: acylated pleckstrin-homology (PH) domain-containing protein; CDPK: Ca^2+^-dependent protein kinase; DAG: diacylglycerol; PA: phosphatidic acid; PKG: protein kinase G.

Chaturvedi et al (1999) explicitly tested for increase of GRA secretion with Ca^2+^ stimulation by supplying exogenous Ca^2+^ to streptolysin O-permeabilized cells. They saw no change in GRA secretion, but it is not known what other cell processes might be perturbed by this permeabilization treatment, and a range of Ca^2+^ concentrations was not tested.

While a reciprocal response of MIC and GRA secretion to Ca^2+^-based signalling is consistently evident, differences in their relative responses depends on the type of treatment, suggesting that different Ca^2+^-signalling events might drive these processes as is also the case for other infection-cycle events (Lourido et al., 2012). For instance, thapsigargin elevates cytosolic Ca^2+^, but while this does lead to microneme secretion, it does not lead to increased motility or conoid extrusion without increased extracellular Ca^2+^ (Pace, Mcknight, Liu, Jimenez, & Moreno, 2014). We also observed no change in GRA secretion with thapsigargin, although MIC secretion does increase. Similarly, cGMP agonists BIPPO and zaprinast both resulted in strong MIC secretion increase, but much more subdued GRA secretion inhibition compared to the Ca^2+^ ionophores. These data indicate that the regulation of microneme exocytosis is more sensitive to Ca^2+^ than the regulation of dense granule exocyotsis. Alternatively, cGMP-based signalling might contribute to some MIC secretion that is independent of Ca^2+^, such as lipid-mediated control of microneme exocytosis (Figure 6) (Bullen et al., 2016).

The comparison of the *Tg*CDPK1 and *Tg*CDPK3 mutants provides further evidence of a separation between the mechanisms of Ca^2+^-mediated control of microneme versus dense granule secretion. Depletion of either of these kinases results in loss of Ca^2+^-dependent inhibition of GRA secretion, suggesting that substrates of both are required for this process. However, the MIC secretion response is quite different between *Tg*CDPK1 and *Tg*CDPK3 depletion: *Tg*CDPK1 loss maintains a Ca^2+^-dependent MIC response although greatly reduced in magnitude; whereas *Tg*CDPK3 loss results in Ca^2+^ insensitivity. This is consistent with some cooperativity between these two kinases where *Tg*CDPK3 might activate the responsiveness of *Tg*CDPK1 for MIC secretion, as others have suggested (Treeck et al., 2014). The complexity of Ca^2+^ signalling targets and their dynamics is further indicated by 3-MB-PP1 treatment either before or after egress, which also result in differences to constitutive MIC secretion as well as GRA control. Post-egress 3-MB-PP1 treatment leaves GRA secretion Ca^2+^-responsive, yet pre-egress treatment blocks this. This suggests that some stable, necessary *Tg*CDPK1 phosphorylation of substrates occurs at the point of egress, and that *Tg*CDPK3 substrates are then sufficient for the responsiveness of dense granules to Ca^2+^.

The location of GRA secretion from *Toxoplasma* tachyzoites has not been unambiguously determined, however it has been suggested that dense granules fuse laterally at the parasite cell surface, rather than apically as micronemes and rhoptries do (Dubremetz et al., 1993). In any case, it is unlikely that suppression of dense granule release is a direct effect of increased microneme secretion and competition for space at the site of secretion. Even if they were to share the same exit point, our data show instances of strongly elevated microneme secretion with no change to dense granule secretion (e.g zaprinast 250 µM, thapsigargin 10 µM).

The mechanism for Ca^2+^-mediated control of GRA secretion is currently unclear. Some GRA proteins, that bear trans membrane domains and are membrane-associated after release into the host cell environment, are known be maintained as soluble high-molecular mass protein aggregates within the dense granules prior to release (Labruyere, Lingnau, Mercier, & Sibley, 1999; Lecordier, Mercier, Sibley, & Cesbron-Delauw, 1999; Sibley et al., 1995). GRA1, a highly abundant GRA with two Ca^2+^ binding-domains, has been speculated to potentially play a role in control of this aggregated state (Lebrun, Carruthers, & Cesbron-Delauw, 2013). It is conceivable that switching from aggregated to disaggregated state of GRAs has a role in secretion regulation and that Ca^2+^ could modulate this. If so, dense granule luminal Ca^2+^ levels would be relevant and must also be controlled. Irrespective of such an internal control process, cytosolic factors are likely to be implicated given evidence of both *Tg*CDPK1 and *Tg*CDPK3 substrates participating in GRA regulation. These substrates might control trafficking of the dense granules or derived vesicles to the relevant location for exocytosis, and/or mediate fusion with the plasma membrane. Several proteins implicated in vesicular trafficking have been identified as CDPK targets in *Plasmodium* (Brochet et al., 2014). If a lateral site of dense granule fusion occurs, some reorganisation of the IMC including membrane cisternae and subpellicular filamentous network would likely be required for vesicular contact with the plasma membrane, and this might require further direction from CDPK1/3-mediated processes.

While the mechanism for Ca^2+^-controlled secretion of dense granules is currently unknown, reciprocal control of micronemes and dense granules using the same signals has a clear biological logic. The protein cargos are mutually exclusive in terms of function—one for the extracellular processes, the other for the intracellular processes. After successful parasite invasion of a host cell, the suppression of MIC secretion by cAMP-mediated depletion of cytosolic Ca^2+^ would concomitantly allow for relaxation of GRA secretion suppression. Furthermore, the coupling of these signalling networks reinforces the potential for signal disruption as a druggable therapeutic strategy. Dysregulation of both MIC and GRA secretion could both interrupt control of the lytic invasion cycle and promote immune-recognition of a wide suite of GRA proteins that might otherwise only be released in the intracellular context.

### Experimental procedures

#### Parasites cultures

*T. gondii* tachyzoites were grown by serial passage in human foreskin fibroblast (HFF) cells as previously described (Striepen B, 2007). Briefly, *Toxoplasma* RH strain parasites were serially passaged in confluent HFF cells containing in ED1 media (Dulbecco’s Modified Eagle’s Medium (DMEM) supplemented with 1% Foetal Bovine Serum (FBS), 0.2mM additional L-Glut, 50 Units/ml Penicillin/Streptomycin, and 0.25 μg/ml of amphotericin-B).

#### Secretion Assays

Parasite cultures were pre-incubated for at least 48 hours with or without ATc. Parasites were harvested after multiple rounds of parasite replication within the parasitophorous vacuole and with approximately 50-80% of vacuoles intact prior to natural egress. Cultures were scraped, host cells disrupted by syringe passing through a 26 guage needle, filtered through a 3 µm polycarbonate filter to remove host debris, and parasites pelleted at 1000x *g* at 15 °C for 10 minutes. Pellets were aspirated and washed with 3 ml of Invasion Buffer (DMEM with 3% FBS and 10 mM HEPES, pH 7.4) and pelleted as before. Supernatants were aspirated and pellets resuspended in Invasion Buffer at 2.5×10^8^ cells.ml^−1^. 50 μl of parasite suspension was then mixed with an equal volume of Invasion Buffer containing 2x the final concentration of agonist or vehicle (DMSO) equivalent. Parasite samples were incubated at 37 °C for 20 minutes to allow secretion and quenched on ice for 2 minutes to stop secretion. Cells were then separated from supernatants by centrifugation (8000 rpm, 2 minutes, 4 °C), 85 μl of supernatant removed and this was centrifuged a second time to remove any remaining cells, with a final volume of 75 µl carefully aspirated. The cell pellets were washed with 1x PBS, repelleted as before, and the supernatant was aspirated. Supernatant and cell pellet proteins were solubilized in SDS Sample Buffer, separated by SDS-PAGE, transferred to nitrocellulose membranes and GRA, MIC or mitochondrial proteins immuno-detected. Western blot detection was performed with Horse Radish Peroxidase conjugated secondary antibodies detected using SuperSignal West Pico Chemiluminescent Substrate (Pierce). Signal strength was quantified using a BioRad Chemidoc imager and ImageLab software, and replicate assays signals were normalised to their respective cell pellet TOM40 signal to control for any minor cell number variation.

## Acknowledgements

We would like to thank Philip Thompson for BIPPO, Chris Tonkin for technical advice and gifting the 3-MB-PP1 inhibitor and *Tg*CDPK3 KO cell line, Sebastian Lourido for gifting the *Tg*CDPK1 iHA KD cell line, Oliver Billker for gifting Compound 2, David Sibley for gifting the MIC2 antibody, Vern Carruthers for gifting the MIC5 antibody and Corinne Mercier for gifting the GRA1, GRA2 and GRA5 antibodies. This work was supported by the Australian Research Council (DP120100599) and the Medical Research Council (UK: MR/M011690/1).

## Reference

Adl, S. M., Simpson, A. G. B., Lane, C. E., Lukes, J., Bass, D., Bowser, S. S., et al. (2012). The revised classification of eukaryotes. Journal of Eukaryotic Microbiology, 59(5), 429–493. http://doi.org/10.1111/j.1550-7408.2012.00644.x

Billker, O., Lourido, S., & Sibley, L. D. (2009). Calcium-dependent signaling and kinases in apicomplexan parasites. Cell Host Microbe, 5(6), 612–622. http://doi.org/10.1016/j.chom.2009.05.017

Blader, I. J., Coleman, B. I., Chen, C.-T., & Gubbels, M.-J. (2015). Lytic cycle of *Toxoplasma gondii*: 15 years later. Annual Review of Microbiology, 69, 463–485. http://doi.org/10.1146/annurev-micro-091014-104100

Borges-Pereira, L., Budu, A., Mcknight, C. A., Moore, C. A., Vella, S. A., Hortua Triana, M. A., et al. (2015). Calcium signaling throughout the *Toxoplasma gondii* lytic cycle: a study using genetically encoded calcium indicators. Journal of Biological Chemistry, 290(45), 26914–26926. http://doi.org/10.1074/jbc.M115.652511

Brochet, M., Collins, M. O., Smith, T. K., Thompson, E., Sebastian, S., Volkmann, K., et al. (2014). Phosphoinositide metabolism links cGMP-dependent protein kinase G to essential Ca2+ signals at key decision points in the life cycle of malaria parasites. PLOS Biology, 12(3), e1001806. http://doi.org/10.1371/journal.pbio.1001806

Brydges, S. D., Harper, J. M., Parussini, F., Coppens, I., & Carruthers, V. B. (2008). A transient forward-targeting element for microneme-regulated secretion in *Toxoplasma gondii*. Biology of the Cell, 100(4), 253–264. http://doi.org/10.1042/BC20070076

Bullen, H. E., Jia, Y., Yamaryo-Botté, Y., Bisio, H., Zhang, O., Jemelin, N. K., et al. (2016). Phosphatidic acid-mediated signaling regulates microneme secretion in *Toxoplasma*. Cell Host and Microbe, 19(3), 349–360. http://doi.org/10.1016/j.chom.2016.02.006

Carruthers, V. B., & Sibley, L. D. (1997). Sequential protein secretion from three distinct organelles of *Toxoplasma gondii* accompanies invasion of human fibroblasts. European Journal of Cell Biology, 73(2), 114–123.

Carruthers, V. B., Giddings, O. K., & Sibley, L. D. (1999a). Secretion of micronemal proteins is associated with toxoplasma invasion of host cells. Cell Microbiol, 1(3), 225–235.

Carruthers, V., Moreno, S., & Sibley, L. (1999b). Ethanol and acetaldehyde elevate intracellular [Ca2+] and stimulate microneme discharge in *Toxoplasma gondii*. The Biochemical Journal, 342, 379–386.

Chaturvedi, S., Qi, H., Coleman, D., Rodriguez, A., Hanson, P. I., Striepen, B., et al. (1999). Constitutive calcium-independent release of *Toxoplasma gondii* dense granules occurs through the NSF/SNAP/SNARE/Rab machinery. The Journal of Biological Chemistry, 274(4), 2424–2431. http://doi.org/10.1074/jbc.274.4.2424

Coppens, I., Andries, M., Liu, J. L., & Cesbron-Delauw, M.-F. (1999). Intracellular trafficking of dense granule proteins in *Toxoplasma gondii* and experimental evidences for a regulated exocytosis. European Journal of Cell Biology, 78, 463–472. http://doi.org/10.1016/S0171-9335(99)80073-9

de Vargas, C., Audic, S., Henry, N., Decelle, J., Mahé, F., Logares, R., et al. (2015). Ocean plankton. Eukaryotic plankton diversity in the sunlit ocean. Science, 348(6237), 1261605. http://doi.org/10.1126/science.1261605

Donald, R. G. K., Allocco, J., Singh, S. B., Nare, B., Salowe, S. P., Wiltsie, J., & Liberator, P. A. (2002). *Toxoplasma gondii* cyclic GMP-dependent kinase: chemotherapeutic targeting of an essential parasite protein kinase. Eukaryotic Cell, 1(3), 317–328. http://doi.org/10.1128/EC.1.3.317-328.2002

Dubremetz, J. F., Achbarou, A., Bermudes, D., & Joiner, K. A. (1993). Kinetics and pattern of organelle exocytosis during *Toxoplasma-gondii* host-cell interaction. Parasitology Research, 79(5), 402–408. http://doi.org/10.1007/BF00931830

Endo, T., Tokuda, H., Yagita, K., & Koyama, T. (1987). Effects of extracellular potassium on acid release and motility initiation in *Toxoplasma gondii*. J. Protozool., 34(3), 291–295.

Etheridge, R. D., Alaganan, A., Tang, K., Lou, H. J., Turk, B. E., & Sibley, L. D. (2014). The *Toxoplasma* pseudokinase ROP5 forms complexes with ROP18 and ROP17 kinases that synergize to control acute virulence in mice. Cell Host and Microbe, 15(5), 537–550. http://doi.org/10.1016/j.chom.2014.04.002

Farrell, A., Thirugnanam, S., Lorestani, A., Dvorin, J. D., Eidell, K. P., Ferguson, D. J. P., et al. (2012). A DOC2 protein identified by mutational profiling is essential for apicomplexan parasite exocytosis. Science, 335(6065), 218–221. http://doi.org/10.1126/science.1210829

Frénal, K., Dubremetz, J.-F., Lebrun, M., & Soldati-Favre, D. (2017). Gliding motility powers invasion and egress in Apicomplexa. Nature Reviews Microbiology, 15(11), 645–660. http://doi.org/10.1038/nrmicro.2017.86

Graindorge, A., Frénal, K., Jacot, D., Salamun, J., Marq, J.-B., & Soldati-Favre, D. (2016). The conoid associated motor MyoH is indispensable for *Toxoplasma gondii* entry and exit from host cells. PLOS Pathogens, 12(1), e1005388. http://doi.org/10.1371/journal.ppat.1005388

Guérin, A., Corrales, R. M., Parker, M. L., Lamarque, M. H., Jacot, D., Hajj, El, H., et al. (2017). Efficient invasion by *Toxoplasma* depends on the subversion of host protein networks. Nature Microbiology, 2(10), 1358–1366. http://doi.org/10.1038/s41564-017-0018-1

Hakimi, M.-A., Olias, P., & Sibley, L. D. (2017). *Toxoplasma* effectors targeting host signaling and transcription. Clinical Microbiology Reviews, 30(3), 615–645. http://doi.org/10.1128/CMR.00005-17

Håkansson, S., Charron, A. J., & Sibley, L. D. (2001). *Toxoplasma* evacuoles: a two-step process of secretion and fusion forms the parasitophorous vacuole., 20(12), 3132–3144. http://doi.org/10.1093/emboj/20.12.3132

Howard, B. L., Harvey, K. L., Stewart, R. J., Azevedo, M. F., Crabb, B. S., Jennings, I. G., et al. (2015). Identification of potent phosphodiesterase inhibitors that demonstrate cyclic nucleotide-dependent functions in apicomplexan parasites. ACS Chemical Biology. http://doi.org/10.1021/cb501004q

Jia, Y., Marq, J.-B., Bisio, H., Jacot, D., Mueller, C., Yu, L., et al. (2017). Crosstalk between PKA and PKG controls pH-dependent host cell egress of *Toxoplasma gondii*. EMBO Journal, 36(21), 3250–3267. http://doi.org/10.15252/embj.201796794

Kafsack, B. F. C., Pena, J. D. O., Coppens, I., Ravindran, S., Boothroyd, J. C., & Carruthers, V. B. (2009). Rapid membrane disruption by a perforin-like protein facilitates parasite exit from host cells. Science, 323(5913), 530–533. http://doi.org/10.1126/science.1165740

Katris, N. J., van Dooren, G. G., McMillan, P. J., Hanssen, E., Tilley, L., & Waller, R. F. (2014). The apical complex provides a regulated gateway for secretion of invasion factors in *Toxoplasma*. PLOS Pathogens, 10(4), e1004074. http://doi.org/10.1371/journal.ppat.1004074

Labruyere, E., Lingnau, M., Mercier, C., & Sibley, L. D. (1999). Differential membrane targeting of the secretory proteins GRA4 and GRA6 within the parasitophorous vacuole formed by *Toxoplasma gondii*. Molecular and Biochemical Parasitology, 102(2), 311–324.

Lebrun, M., Carruthers, V. B., & Cesbron-Delauw, M.-F. (2013). *Toxoplasma gondii* The Model Apicomplexan - Perspectives and Methods. In L. M. Weiss & K. Kim (Eds.), Toxoplasma secretory proteins and their roles in cell invasion and intracellular survival (Second Edition, pp. 389–453). Elsevier. http://doi.org/10.1016/B978-0-12-396481-6.00012-X

Lecordier, L., Mercier, C., Sibley, L. D., & Cesbron-Delauw, M. F. (1999). Transmembrane insertion of the *Toxoplasma gondii* GRA5 protein occurs after soluble secretion into the host cell. Molecular Biology of the Cell, 10(4), 1277–1287.

Lourido, S., Jeschke, G. R., Turk, B. E., & Sibley, L. D. (2013). Exploiting the unique ATP-binding pocket of *Toxoplasma* calcium-dependent protein kinase 1 to identify its substrates. ACS Chemical Biology, 8(6), 1155–1162. http://doi.org/10.1021/cb400115y

Lourido, S., Shuman, J., Zhang, C., Shokat, K. M., Hui, R., & Sibley, L. D. (2010). Calcium-dependent protein kinase 1 is an essential regulator of exocytosis in *Toxoplasma*. Nature, 465(7296), 359–362. http://doi.org/10.1038/nature09022

Lourido, S., Tang, K., & Sibley, L. D. (2012). Distinct signalling pathways control *Toxoplasma* egress and host-cell invasion. Embo J, 31(24), 4524–4534. http://doi.org/10.1038/emboj.2012.299

Mahé, F., de Vargas, C., Bass, D., Czech, L., Stamatakis, A., Lara, E., et al. (2017). Parasites dominate hyperdiverse soil protist communities in Neotropical rainforests. Nature Ecology & Evolution, 1(4), 91. http://doi.org/10.1038/s41559-017-0091

McCoy, J. M., Whitehead, L., van Dooren, G. G., & Tonkin, C. J. (2012). *Tg*CDPK3 regulates calcium-dependent egress of *Toxoplasma gondii* from host cells. PLOS Pathog, 8(12), e1003066. http://doi.org/10.1371/journal.ppat.1003066.s010

Mercier, C., & Cesbron-Delauw, M.-F. (2015). *Toxoplasma* secretory granules: one population or more? Trends in Parasitology, 31(2), 60–71. http://doi.org/10.1016/j.pt.2014.12.002

Mercier, C., Adjogble, K. D. Z., Däubener, W., & Delauw, M.-F.-C. (2005). Dense granules: are they key organelles to help understand the parasitophorous vacuole of all apicomplexa parasites? International Journal for Parasitology, 35(8), 829–849. http://doi.org/10.1016/j.ijpara.2005.03.011

Pace, D. A., Mcknight, C. A., Liu, J., Jimenez, V., & Moreno, S. N. J. (2014). Calcium entry in *Toxoplasma gondii* and its enhancing effect of invasion-linked traits. Journal of Biological Chemistry. http://doi.org/10.1074/jbc.M114.565390

Paul, A. S., Saha, S., Engelberg, K., Jiang, R. H. Y., Coleman, B. I., Kosber, A. L., et al. (2015). Parasite calcineurin regulates host cell recognition and attachment by apicomplexans. Cell Host and Microbe, 18, 1–12. http://doi.org/10.1016/j.chom.2015.06.003

Schlossmann, J., Ammendola, A., Ashman, K., Zong, X., Huber, A., Neubauer, G., et al. (2000). Regulation of intracellular calcium by a signalling complex of IRAG, IP3 receptor and cGMP kinase Ibeta. Nature, 404(6774), 197–201. http://doi.org/10.1038/35004606

Sibley, L. D., Niesman, I. R., Parmley, S. F., & Cesbron-Delauw, M. F. (1995). Regulated secretion of multi-lamellar vesicles leads to formation of a tubulo-vesicular network in host-cell vacuoles occupied by *Toxoplasma gondii*. Journal of Cell Science, 108 (Pt 4), 1669–1677.

Sidik, S. M., Hortua Triana, M. A., Paul, A. S., Bakkouri, El, M., Hackett, C. G., Tran, F., et al. (2016). Using a Genetically Encoded Sensor to Identify Inhibitors of *Toxoplasma gondii* Ca2+ Signaling. Journal of Biological Chemistry, 291(18), 9566–9580. http://doi.org/10.1074/jbc.M115.703546

Stewart, R. J., Whitehead, L., Nijagal, B., Sleebs, B. E., Lessene, G., McConville, M. J., et al. (2017). Analysis of Ca2+ mediated signaling regulating *Toxoplasma* infectivity reveals complex relationships between key molecules. Cellular Microbiology, 19(4). http://doi.org/10.1111/cmi.12685

Tang, Q., Andenmatten, N., Hortua Triana, M. A., Deng, B., Meissner, M., Moreno, S. N. J., et al. (2014). Calcium-dependent phosphorylation alters class XIVa myosin function in the protozoan parasite *Toxoplasma gondii*. Molecular Biology of the Cell, 25(17), 2579–2591. http://doi.org/10.1091/mbc.E13-11-0648

Treeck, M., Sanders, J. L., Gaji, R. Y., LaFavers, K. A., Child, M. A., Arrizabalaga, G., et al. (2014). The calcium-dependent protein kinase 3 of *Toxoplasma* influences basal calcium levels and functions beyond egress as revealed by quantitative phosphoproteome analysis. PLOS Pathogens, 10(6), e1004197. http://doi.org/10.1371/journal.ppat.1004197

Uboldi, A. D., Wilde, M.-L., McRae, E. A., Steward, R. J., Dagley, L. F., Yang, L., et al. (2018). Protein Kinase A Negatively Regulates Ca2+ signalling in *Toxoplasma gondii*. PLOS Biology (in press)

Wetzel, D. M., Chen, L. A., Ruiz, F. A., Moreno, S. N. J., & Sibley, L. D. (2004). Calcium-mediated protein secretion potentiates motility in *Toxoplasma gondii*. Journal of Cell Science, 117(Pt 24), 5739–5748. http://doi.org/10.1242/jcs.01495

Wiersma, H. I., Galuska, S. E., Tomley, F. M., Sibley, L. D., Liberator, P. A., & Donald, R. G. K. (2004). A role for coccidian cGMP-dependent protein kinase in motility and invasion. International Journal for Parasitology, 34(3), 369–380. http://doi.org/10.1016/j.ijpara.2003.11.019

